# Genotypic and Phenotypic Analyses of Two Distinct Sets of *Pseudomonas aeruginosa* Urinary Tract Isolates

**DOI:** 10.1101/2023.12.21.572880

**Authors:** HA Ebrahim, S Haldenby, MP Moore, AA Dashti, RV Floyd, JL Fothergill

## Abstract

Urinary tract infections (UTIs) are associated with a high burden of morbidity, mortality, and cost. *Pseudomonas aeruginosa* employs a myriad of virulence factors, including biofilm formation and motility mechanisms, to cause infections including persistent UTIs. *P. aeruginosa* is highly resistant to antibiotics and the World Health Organization has identified it as a pathogen for which novel antimicrobials are urgently required. Genotypic and phenotypic characterization of *P. aeruginosa* from UTIs are underreported. In addition, the rise of antimicrobial resistance (AMR) is a cause for concern, particularly in many countries where surveillance is severely lacking.

22 *P. aeruginosa* UTI isolates were sourced from the United Kingdom (UK) and Kuwait. To establish the phenotypes of UK isolates, growth analysis, biofilm formation assays, motility assays, and antibiotic disc diffusion assays were performed. Whole genome sequencing, antimicrobial susceptibility assays, and *in silico* detection of AMR-associated genes were conducted on both sets of isolates.

In terms of their phenotypic characteristics and genomic composition, the UTI isolates varied. Multiple resistance genes associated with resistance to various classes of antibiotics, such as aminoglycosides, fluoroquinolones, and β-lactams, particularly in isolates from Kuwait. Extreme antibiotic resistance was detected in the isolates obtained from Kuwait, indicating that the country may be an antibiotic resistance hotspot.

This study highlights that isolates from UTIs are diverse and can display extremely high resistance. Surveillance in countries such as Kuwait are currently limited and this study suggest the need for greater surveillance.

## Introduction

Urinary tract infections (UTIs) are some of the most widespread infections in healthcare and community settings worldwide, with around 150 million patients infected annually (1,2). According to the National Health Service in England, UTIs resulted in more than 800,000 hospital visits in the period between 2018 and 2023 (3). UTIs are also one of the main causes of serious *Escherichia coli* (*E. coli*) bloodstream infections and a major contributor to the rise of antimicrobial resistance (AMR) in England (4).

*Pseudomonas aeruginosa* is a Gram-negative, motile, facultative anaerobic bacteria (5). *P. aeruginosa* cause opportunistic infections in people with cystic fibrosis, cancer, burns, urinary tract complications and chronic wounds (6). The World Health Organization (WHO) declared *P. aeruginosa* as a microorganism of concern and in urgent need of new medical interventions (7). Nosocomial infections caused by *P. aeruginosa* are often difficult to treat with antibiotics due to intrinsic and acquired multi-drug resistance (8). *P. aeruginosa* also utilises biofilm formation, which can complicate treatment in respiratory, urinary tract and eye infections (9,10). The bacterium utilises a complex system of regulatory circuits via quorum sensing, two component systems and alternative sigma factors along with other regulators (11). The large complex genome of *P. aeruginosa* (6-7 MB) contributes to adaptation in different environments and niches (12). In addition*, P. aeruginosa* utilises multiple virulence factors such as phenazines, elastase, phospholipase C, alkaline protease, rhamnolipids and hydrogen cyanide (13). In a murine model of catheter associated infections (CAUTI), *P. aeruginosa* utilises type III secretion system to cause acute infections (14).

*P. aeruginosa* is efficient in causing acute or recurrent UTI, partially due to its ability to form biofilms and survive intracellularly (15). The primary route to the urinary tract occurs in nosocomial settings, UTIs account for 40% of hospital acquired infections (5). The percentage of CAUTIs attributed to *P. aeruginosa* is estimated at 35%, indicating that the pathogen is proficient at causing these infections (5). The presence of patients in hospitals for more than 30 days increases the risk of acquiring UTIs by almost 100% (16). Infections caused by CAUTI *P. aeruginosa* are usually known to be severe, persistent and antibiotic resistant (17).

While surveillance of the prevalence antimicrobial resistance of UTI *P. aeruginosa* in the United Kingdom occurs, it is not comprehensive. Ironmonger *et al.* (2015) conducted a comprehensive antimicrobial resistance surveillance in the Midlands region of the United Kingdom to identify resistance patterns of uropathogens, including *P. aeruginosa*. A total of 786 *Pseudomonas* spp isolates (4.08% of all uropathogens) acquired over a four-year period from a population of 5.6 million, 5.7 % carbapenemase producers were identified (18). However, many other phenotypic features are not reported. Furthermore, the similarity of these data to other countries worldwide is unclear. Minimal reporting on UTIs is common. Including in the Gulf Corporation Council (GCC), which is comprised of 6 countries: The state of Kuwait, Kingdom of Saudi Arabia, Kingdom of Bahrain, State of Qatar, United Arab Emirates and Sultanate of Oman. Current evidence suggests that *P. aeruginosa* resistance to antibiotics is rapidly evolving in the GCC countries (19,20). The increasing rates of antimicrobial resistance are attributed to several factors such as travel (21,22), high use of antibiotics (23,24) including antibiotics sales from pharmacies without prescription (19,25). Zowawi *et al* conducted a regional study *on P. aeruginosa* clinical isolates and found that high risk clones were widespread, in particular *bla*_VIM-_ was detected in 39% of a total of 95 isolates (20). Consistent surveillance on the spread of *P. aeruginosa* within Kuwaiti hospital is lacking. Multiple studies reveal the existence of multidrug resistance *P. aeruginosa* in nosocomial settings. In 1997, a study reported 10 UTI *P. aeruginosa* isolates with aminoglycosides being the most common resisted class of antibiotics. A study conducted over a three-year period (2005–2007) in one of the largest hospitals in Kuwait revealed that 9.8 percent of *P. aeruginosa* UTIs were acquired in the hospital, compared to 4.33% isolated from outpatients, 15% and 14% of hospital-acquired isolates were resistant to amikacin and piperacillin/tazobactam, respectively (26). However, these published reports lack detailed genomic analyses on the carriage of AMR genes.

In this study, we highlight genotypic and phenotypic characteristics of *P. aeruginosa* UTI isolates obtained in the United Kingdom and compare to a small subset of isolates from the State of Kuwait.

## Methods

### Bacterial isolates

*P. aeruginosa* isolates were obtained from the Royal Liverpool University Hospital in Liverpool (n=15) and the State of Kuwait (n=8). In the UK panel, the average age of patients on upon bacterial isolation was 70 years old with 47% isolates from woman. All isolates sourced from Kuwait were isolated in 2005 with no further information available.

### Isolate storage and culture

The isolates were stored at −80 °C in Luria-Bertani (LB) broth (Sigma Aldrich) (W/V) 10% glycerol (Sigma Aldrich). Isolates were streaked onto LB agar (Appleton Woods) and grown overnight at 37 °C. A single colony of each *P. aeruginosa* isolate was utilized to inoculate LB broth (Sigma Aldrich) in universal tubes overnight. Subsequently, the cultures were grown at 37°C and shaking at 180 rpm.

### Growth analysis

To establish the rate of bacterial growth. Overnight cultures were diluted 1:100 in polystyrene 96-well plates (Corning® Costar®). The assay was performed with four technical replicates and in quadruplicate biological replicates. The growth of each well was monitored for 24 h at 37°C and the absorbance 600nm was recorded every 30 min by the Fluostar Omega plate reader. Analysis was performed using Growthcurver in R Studio.

### Initial biofilm attachment assay

The overnight cultures of the UK based clinical isolates were diluted 1:100 in LB. This was followed by the addition of 200µl of each *P. aeruginosa* containing-LB solution in quadruplicate to a 96-well plate (Corning® Costar®) and grown at 37°C for 24h or 48h. After the incubation period, the broth was removed, followed by the application of 200µl of phosphate buffered saline (PBS) to the wells twice to wash the biofilms. Removal of PBS was conducted followed by drying at 37 C. 200µl of 0.25% crystal violet (CV) w/v in dH2o (Sigma Aldrich) was added into each well for 10 minutes. Stained biofilms were solubilised by adding 200µl of 95% ethanol v/v (Sigma Aldrich) for 10 minutes and transferred to a new 96-well plate (Corning® Costar®). The absorbance was measured at OD600nm on the Fluorostar Omega plate reader to determine the biofilm biomass.

### Motility Assays

#### Twitching

To measure twitching motility of the isolates, a colony of *P. aeruginosa* was stabbed in the middle of an agar plate to the bottom. The colonies were then grown overnight at 37°C. The agar was then removed, and the plate was subsequently stained with 0.25 % CV for 15 minutes. Twitching zones were measured using a ruler. The cut-off point in which the isolate was considered motile was 5 mm in diameter.

#### Swimming motility

The methodology used is based on a swimming motility assay by Ha, *et al,* (2014) (30). Briefly, media was prepared according to the protocol described. *P. aeruginosa* cultures were grown overnight in LB broth at 37 °C shaking at 180 rpm. Subsequently, a sterile toothpick was used to stab the agar from top to near bottom of the plate. The agar plates were incubated at 37 °C for 18 h. Three biological replicates were performed for each isolate in the study. Millimeter measurements were taken to assess the diameter of swimming bacteria. To determine whether the isolate is motile, comparison with the reference laboratory strain PA14 were conducted.

#### Swarming motility

The swarming motility test was performed according to the protocol established by Ha, *et al,* (2014) (30). Once the preparations of plate cultures were completed, 2.5 µl of overnight of *P. aeruginosa* cultures were added to the top of the agar. The agar plates were incubated overnight at 37 °C for 18 h. 3 biological replicates were conducted for each isolate. Measurements in mm were taken and the isolates were deemed motile/non-motile depending on the diameter of 6mm.

### Antibiotic resistance and genomic sequencing

#### Disc diffusion assay

Antibiotic susceptibility testing was performed based on the European Committee on Antimicrobial Susceptibility Testing (EUCAST) protocol (31). Three biological replicates of each clinical isolate were grown overnight on Mueller-Hinton agar with antibiotics disc (Thermo Fisher Scientific). The measurements in mm were determined and calculated as an average of 3 readings and compared to the published EUCAST breakpoints to determine if the clinical isolate is sensitive, increased exposure or resistant.

#### Minimum inhibitory concentrations of antimicrobial agents

Minimum inhibitory concentrations (MIC) for 1 clinical UTI *P. aeruginosa* isolate 758 sourced from Kuwait, was determined using the microdilution method as described in the guidelines of EUCAST (32). Briefly, 100µl of overnight culture of the isolate was diluted to OD600 of 0.05 and was mixed with 100µl of serially diluted colisitin (512 - 0.5 μg/ml) (Sigma, UK) in triplicate using cation adjusted MH broth. Microtitre plates were incubated for 1 - 2 days at 37 °C without shaking and bacterial growth was determined by measuring absorbance at OD600 with a FLUOstar® Omega microplate reader and the MARS Data Analysis Software test.

#### Genomic DNA analysis

After extraction with the Wizard® Genomic DNA Purification Kit (Promega), all isolates in this study were sequenced. The DNA was preserved between 2 and 8 degrees Celsius in preparation for sequencing at the University of Liverpool’s Centre for Genomic Research (CGR). Specifically, two isolates were sequenced for each panel sample. Using an Illumina HiSeq 2000 sequencer, 100 bp paired reads were produced from the ends of 500 bp fragments. Upon detection of Illumina adapter sequences using Cutdapt version 1.2.1 (33), FastQ files containing sequenced read data were truncated. This was followed by the selection of option −O 3, which resulted in the 3’ ends of all reads matching the adapter sequence by at least 3 bp being trimmed. The Sickle version 1.2 was utilized, and a minimum window quality of 20 was selected as the score for additional sequence trimming (34).

#### Identification of antibiotic resistance genes

The Comprehensive Antibiotic Resistance Database (CARD; http://arpcard.mcmaster.ca) was used to look for resistance genes on the CGR-obtained sequences (35). Prior to analysis, the sequences were entered into the Resistance Gene Identifier (RGI) search tool, and the following options were checked: DNA Sequence, perfect and strict hits only, exclude nudge, and High quality/coverage.

#### Statistical analysis

All statistical analyses (unless otherwise specified) were conducted using Prism 9.5.1 and R Studio. The Shapiro-Wilk test was used to assess the distribution. Biofilm assays and growth curves were analysed with Kruskal-Wallis analysis of variance, Holm-Sidak post hoc test for nonparametric data, and Brown-Forsyth test for pairwise comparison. Kruskal-Wallis analysis of variance and the Brown-Forsythe test were utilized to analyse the MIC data.

## Results

### Growth dynamics of *P. aeruginosa* UTI isolates in nutrient rich media

Growth in LB media was monitored for 24 h for the reference strains and the 14 UK-based UTI isolates. The area under the curve was lower than PAO1 for all isolates and for 10/14 UTI isolates, this was significantly reduced (Figure 1). However, the UTI isolates exhibited a high degree of variability in the generation time and overall growth rate. Two UTI isolates had a significantly slower generation time (133044 and 133126) and two displayed a significantly faster generation time (133042 and 133098). Five UTI isolates had a significantly higher growth rate while two isolates were significantly reduced (Figure 1). This highlights the inter-isolate variation. However, as PAO1 is considered well adapted to the laboratory environment, it was notable that many of the UTI displayed similar or faster growth under these conditions.

**Figure 1:**
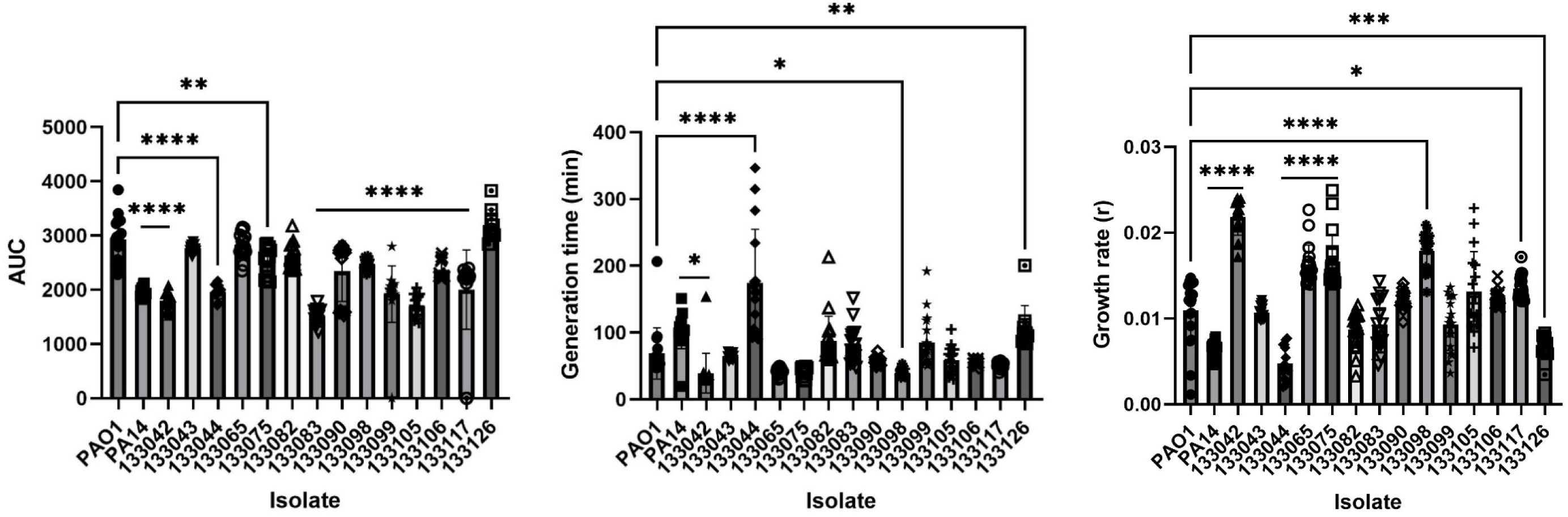
Growth of UTI isolates and 2 references strains (PAO1 and PA14) over 24 h (n≥10). Optical density was measured at OD600 and analysed using Growthcurver to produce A) the area under the curve (AUC), B) generation time and C) growth rate. An ANOVA and Dunnett’s multiple comparison tests were performed between each isolate compared to PAO1. Graphs show individual data points, mean and standard deviation, analysed with a one-way ANOVA and Dunnett’s multiple comparisons test, *p<0.05, **p<0.01, ***p<0.001, ****p<0.0001.

### Biofilm Formation

In order to compare biofilm capability, initial attachment on polystyrene wells was quantified using crystal violet staining. Twelve of fourteen UTI isolates displayed a significant reduction in biofilm compared to the reference strain PAO1, with isolate 13044 showing the lowest biomass (P<0.0001) (Figure 2). The results show that though all isolates showed evidence of biofilm formation, many were significantly reduced compared to PAO1.

**Figure 2:**
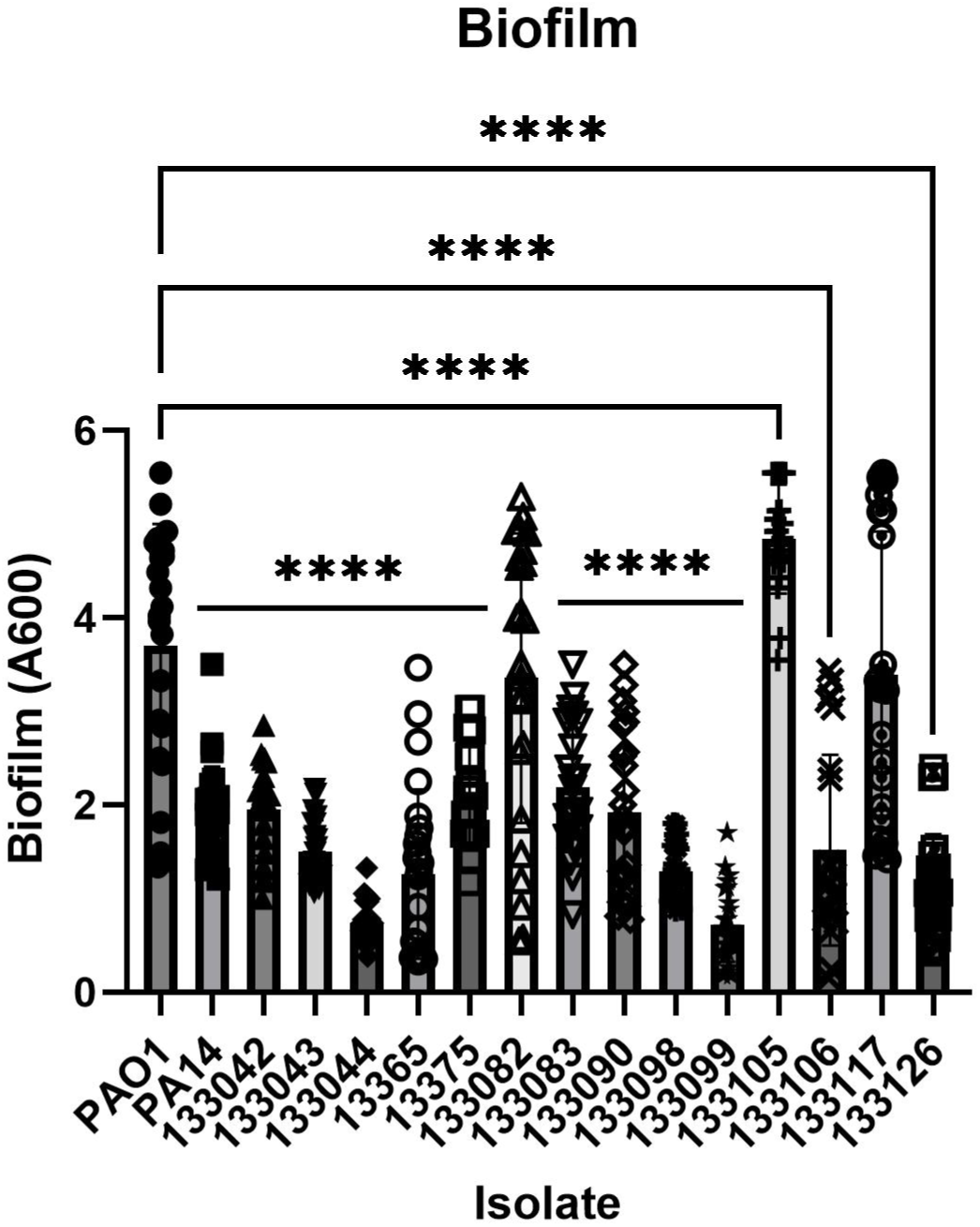
Biofilm formation in the UK-based isolates and reference strains. An ANOVA and Dunnett’s multiple comparison tests were performed between each isolate compared to PAO1. Individual data points, mean and standard deviation, analysed with a one-way ANOVA and Dunnett’s multiple comparisons test are shown. *p<0.05, **p<0.01, ***p<0.001, ****p<0.0001.

### Bacterial motility of UTI isolates

#### Swimming motility

Swimming motility is conducted by a rotating polar flagella and flagellae are also considered a virulence factor which promotes biofilm formation (36). A plate-based method was utilised to assess the swimming ability of the 15 UK-based isolates (30). Upon comparison with the reference strain PAO1 and PA14 and using a cut-off value, 80% of the isolates shown the ability swim through aqueous medium (Figure 3). Isolates 133044,133083 and 13099 were classed as non-motile they were beneath the cut-off value of 6mm and showed a significant reduction in motility compared to PA01.

**Figure 3.**
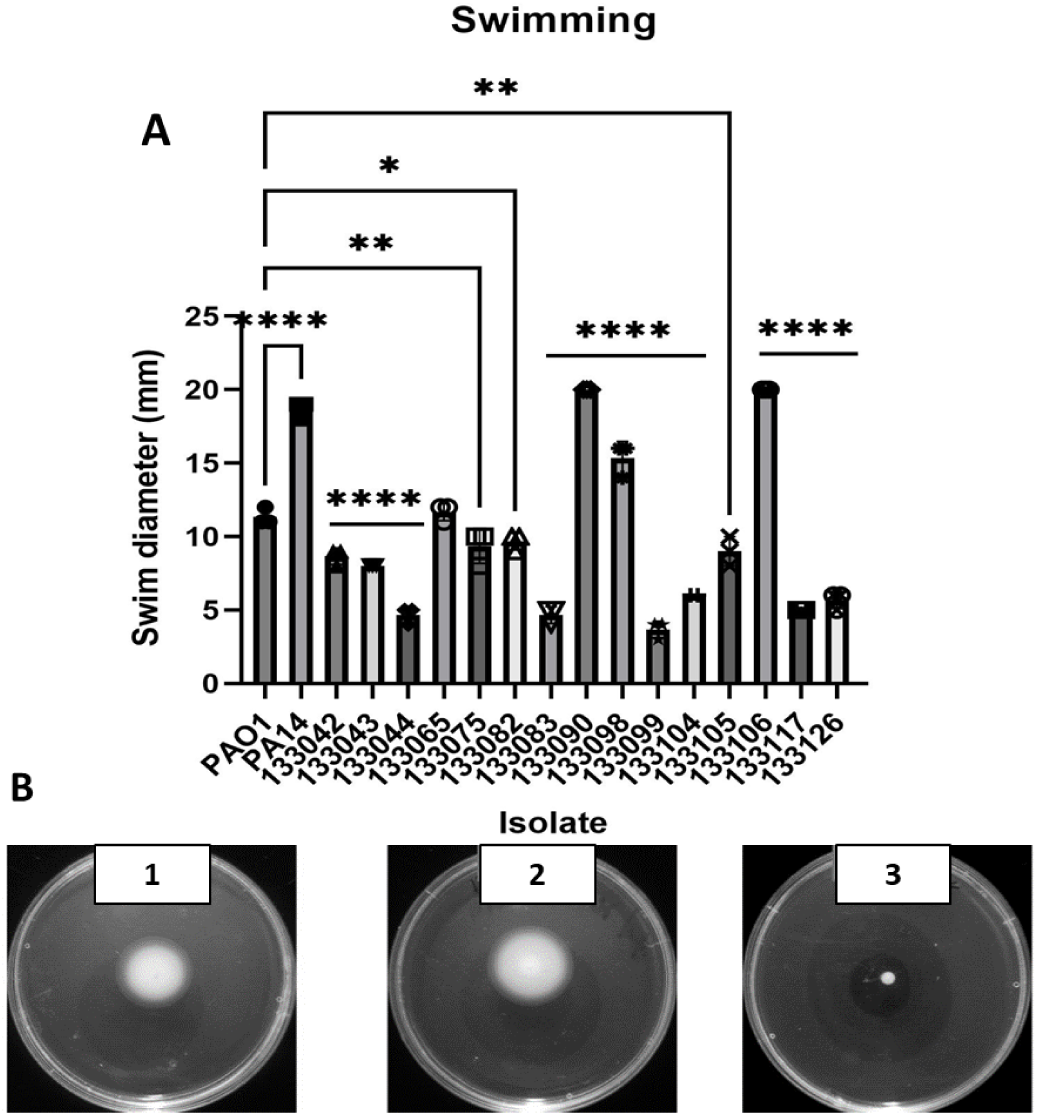
Swimming of *P. aeruginosa U*TI isolates. A) Distances (mm) travelled by each isolate. An ANOVA and Dunnett’s multiple comparison tests were performed between each isolate compared to PAO1. Individual data points, mean and standard deviation, analysed with a one-way ANOVA and Dunnett’s multiple comparisons test are shown. *p<0.05, **p<0.01, ***p<0.001, ****p<0.0001.**B)** The images depict the swimming motility of PA14(1), isolate 133106 (2) and isolate 133099 (image 3).

#### Twitching motility

In contrast to swimming motility, twitching motility is conducted by type IV pili. The pili facilitates movement of *P. aeruginosa* across surfaces, aids colonisation of the host and biofilm formation (37). The mean distance travelled by all the 15 clinical isolates was 6 mm SD=+/−0.75 mm (Figure 4A). The cut-off (5mm) was exceeded by all the UTI isolates except 133044 and 133105 which were non-motile (1 mm). Therefore, the percentage of the isolates capable of performing twitching motility is 86.6%.

**Figure 4:**
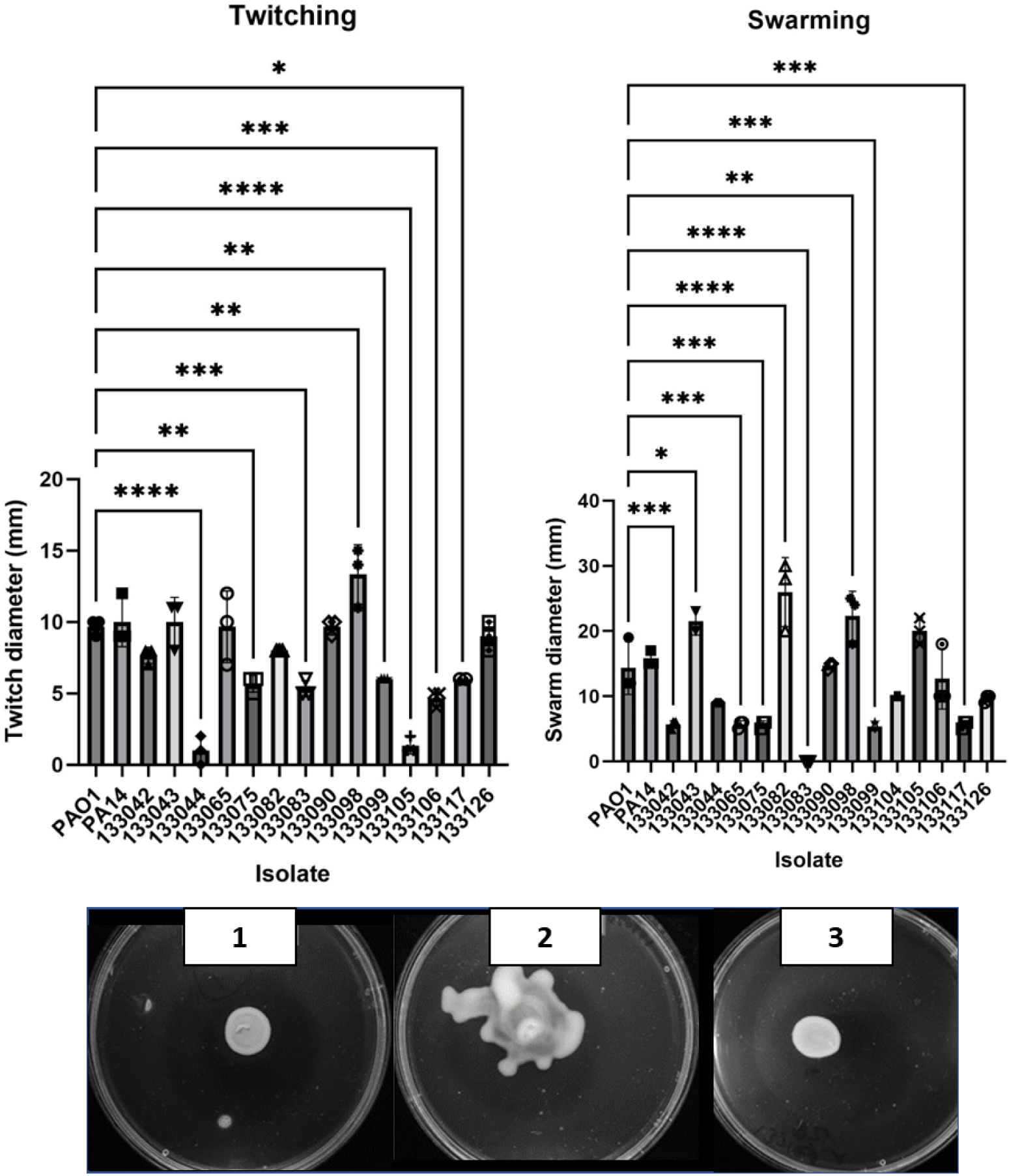
A) Twitching motility. The cut-ff for twitching was 5 mm cut-off motility. B) Swarming motility, the mean average of distance travelled (mm) by swarming motility. The cut-off value was 6 mm. An ANOVA and Dunnett’s multiple comparison tests were performed between each isolate compared to PAO1. Individual data points, mean and standard deviation, analysed with a one-way ANOVA and Dunnett’s multiple comparisons test are shown. *p<0.05, **p<0.01, ***p<0.001, ****p<0.0001. C) Plate images of swarming motility 1: PAO1 2. Isolate 133098 3: Isolate 133082 with moderate swarming ability.

#### Swarming motility

Swarming motility is utilised by *P. aeruginosa* to achieve fast movement across semi-fluid surfaces (38). The cut-off point used to define the motile and the non-motile *P. aeruginosa* was 6 mm (Figure 4B and 4C). The mean distance travelled by all isolates was 6 mm. With the exception of two isolates (133083 and 133099), all UK-based UTI isolates were capable of swarming. Three isolates (133043, 133082 and 133098) were significantly increased in swarming compared to PAO1.

### Antibiotic susceptibility in clinical UTI isolates

In order to test the susceptibility of *P. aeruginosa* to clinical therapeutics, an extensive panel of antibiotics were utilised to examine all clinical isolates and reference isolates PAO1 and PA14 according to clinical breaking points set by EUCAST (39). Table 2 displays the susceptibility testing of the UK isolates. Overall, one isolate (133044) displayed multidrug resistance (defined by resistance to one antibiotic in 3 or more classes). All UK isolates were susceptible with increased exposure to penicillins. For cephalosporins, all isolates were susceptible with increased exposure to the ceftazidime and cefepime, and fully susceptible to ceftazidime/avibactam. One isolate (133044) was resistant to tobramycin, with evidence of resistance to other aminoglycosides but no breakpoints are defined for these. No zones of inhibition were observed when 133044 was exposed to tobramycin, netilmicin or gentamicin. Carbapenems are a key, last resort antibiotic for many *P. aeruginosa* infections. One UK UTI clinical isolates (133044) displayed resistance to imipenem however, the majority of isolates were fully susceptible to meropenem. One isolate (133105) was resistant to aztreonam, with no zone of inhibition (Table 2).

**Table 1:** *P. aeruginosa* isolates used in this study.

| Isolate Name | Source | Date | Country of origin | Gender | Age | Reference |
| --- | --- | --- | --- | --- | --- | --- |
| PAO1 | Wound | 1954 | Australia | N/A | N/A | (27) |
| PA14 | Burn Wound | 1977 | US | N/A | N/A | (28) |
| LESB58 | CF | 1988 | UK- Liverpool | N/A | N/A | (29) |
| 133042 | Urine | 11-Oct-13 | UK | M | 52 | This study |
| 133043 | Urine | 16-Oct-13 | UK | F | 80 | This study |
| 133044 | Urine | 16-Oct-13 | UK | F | 69 | This study |
| 133065 | Urine | 18-Oct-13 | UK | F | 70 | This study |
| 133075 | Urine | 24-Oct-13 | UK | M | 76 | This study |
| 133082 | Urine | 25-Oct-13 | UK | M | 89 | This study |
| 133083 | Urine | 25-Oct-13 | UK | F | 68 | This study |
| 133090 | Urine | 25-Oct-13 | UK | M | 83 | This study |
| 133098 | Urine | 31-Oct-13 | UK | M | 58 | This study |
| 133099 | Urine | 31-Oct-13 | UK | F | 61 | This study |
| 133104 | Urine | 01-Nov-13 | UK | M | 72 | This study |
| 133105 | Urine | 01-Nov-13 | UK | M | 50 | This study |
| 133106 | Urine | 02-Nov-13 | UK | M | 83 | This study |
| 133117 | Urine | 02-Nov-13 | UK | F | 61 | This study |
| 133126 | Urine | 07-Nov-13 | UK | F | 75 | This study |
| 758 | Urine | 4-Jan-05 | Kuwait | N/A | N/A | This study |
| 783 | Urine | 10-Jan-05 | Kuwait | N/A | N/A | This study |
| 786 | Urine | 11-Jan-05 | Kuwait | N/A | N/A | This study |
| 864 | Urine | 18-Apr-05 | Kuwait | N/A | N/A | This study |
| 888 | Urine | 15-May-05 | Kuwait | N/A | N/A | This study |
| 902 | Urine | 22-May-05 | Kuwait | N/A | N/A | This study |
| 925 | Urine | 05-June-05 | Kuwait | N/A | N/A | This study |
| 1083 | Urine | 12-Nov-05 | Kuwait | N/A | N/A | This study |

**Table 2:** Antimicrobial susceptibility of *P. aeruginosa* isolates from UTIs in the UK and Kuwait. The size in mm refers to the zone of inhibition following disk diffusion assays. Clinical breakpoints for resistance (R) are shown in bold, underlined; Susceptible in italics and grey isolates classed as Susceptible, Increased Exposure (I) are shown in black values only.

|  | Tobramycin | Gentamicin | Amikacin | Netilmicin | Imipenem | Meropenem | Doripenem | Ceftazidime | Cefepime | Ceftazidime/avibactam | Levofloxacin | Ciprofloxacin | Aztreonam | Piperacillin | Piperacillin/ Tazobactam | Ticarcillin | Ticarcillin/Clavulanic acid |
| --- | --- | --- | --- | --- | --- | --- | --- | --- | --- | --- | --- | --- | --- | --- | --- | --- | --- |
| UK |  |  |  |  |  |  |  |  |  |  |  |  |  |  |  |  |  |
| 133042 | 24 | 21 | 26 | 18 | 29 | 34 | 35 | 29 | 28 | 28 | 35 | 26 | 30 | 27 | 26 | 26 | 28 |
| 133043 | 24 | 23 | 21 | 17 | 25 | 27 | 35 | 23 | 30 | 22 | 32 | 24 | 20 | 22 | 24 | 25 | 20 |
| 133044 | <u>0</u> | <u>0</u> | 18 | <u>0</u> | 18 | 28 | 31 | 27 | 22 | 27 | 29 | 24 | 22 | 18 | 25 | <u>0</u> | 26 |
| 133065 | 26 | 25 | 29 | 22 | 31 | 31 | 40 | 26 | 28 | 27 | <u>21</u> | <u>12</u> | 22 | 28 | 27 | 33 | 27 |
| 133075 | 28 | 28 | 31 | 22 | 25 | 33 | 32 | 25 | 32 | 25 | 37 | 30 | 19 | 24 | 27 | 23 | 21 |
| 133082 | 23 | 24 | 27 | 18 | 27 | 26 | 28 | 25 | 25 | 25 | 35 | 28 | 21 | 20 | 27 | 24 | 21 |
| 133083 | 22 | 22 | 22 | 19 | 26 | 37 | 35 | 25 | 29 | 25 | 33 | 23 | 26 | 22 | 24 | 28 | 21 |
| 133090 | 25 | 23 | 25 | 19 | 27 | 29 | 31 | 26 | 32 | 28 | 34 | 25 | 27 | 22 | 29 | 25 | 22 |
| 133098 | 26 | 26 | 27 | 22 | 27 | 38 | 32 | 26 | 30 | 26 | 33 | 30 | 29 | 28 | 26 | 28 | 24 |
| 133099 | 21 | 18 | 20 | 12 | 29 | 28 | 28 | 24 | 24 | 25 | <u>0</u> | <u>0</u> | 20 | <u>18</u> | 26 | 23 | 19 |
| 133104 | 21 | 18 | 20 | 12 | 29 | 28 | 28 | 24 | 24 | 25 | <u>0</u> | <u>0</u> | 24 | <u>18</u> | 26 | 23 | 19 |
| 133105 | 21 | 18 | 20 | <u>4</u> | 27 | 28 | 28 | 22 | 22 | 22 | 32 | <u>0</u> | <u>0</u> | 19 | 23 | 22 | 26 |
| 133106 | 25 | 25 | 21 | 20 | 24 | 34 | 34 | 23 | 30 | 23 | 33 | 23 | 25 | 24 | 27 | 21 | 23 |
| 133117 | 20 | 20 | 19 | 10 | 28 | 20 | 30 | 24 | 23 | 21 | <u>0</u> | <u>0</u> | 25 | 20 | 26 | 22 | 21 |
| 133126 | 22 | 19 | 21 | 16 | 27 | 36 | 35 | 24 | 31 | 25 | 26 | 26 | 25 | 24 | 27 | 27 | 25 |
| KUWAIT |  |  |  |  |  |  |  |  |  |  |  |  |  |  |  |  |  |
| 758 | <u>0</u> | <u>11</u> | <u>0</u> | <u>0</u> | <u>9</u> | <u>0</u> | <u>0</u> | <u>0</u> | <u>6</u> | <u>0</u> | <u>0</u> | <u>0</u> | <u>10</u> | <u>0</u> | <u>0</u> | <u>0</u> | <u>0</u> |
| 783 | <u>0</u> | <u>10</u> | <u>0</u> | <u>0</u> | <u>10</u> | <u>0</u> | <u>0</u> | <u>0</u> | <u>5</u> | <u>0</u> | <u>0</u> | <u>0</u> | <u>10</u> | <u>0</u> | <u>0</u> | <u>0</u> | <u>0</u> |
| 786 | <u>0</u> | <u>0</u> | <u>14</u> | <u>3</u> | 24 | <u>16</u> | <u>16</u> | <u>10</u> | <u>11</u> | <u>11</u> | <u>0</u> | <u>0</u> | <u>3</u> | <u>9</u> | <u>12</u> | <u>0</u> | <u>0</u> |
| 864 | 20 | 20 | 25 | 17 | <u>11</u> | <u>0</u> | <u>15</u> | <u>15</u> | <u>20</u> | <u>16</u> | 25 | <u>12</u> | <u>16</u> | <u>14</u> | <u>14</u> | <u>11</u> | <u>8</u> |
| 888 | <u>0</u> | <u>0</u> | <u>0</u> | <u>0</u> | <u>0</u> | <u>11</u> | <u>14</u> | 23 | 24 | 23 | <u>0</u> | <u>0</u> | 18 | <u>7</u> | 19 | <u>14</u> | <u>7</u> |
| 902 | *20* | *20* | *25* | *17* | *25* | *20* | <u>**21**</u> | <u>**10**</u> | <u>**16**</u> | <u>**8**</u> | <u>**22**</u> | <u>**11**</u> | <u>**9**</u> | <u>**0**</u> | <u>**9**</u> | <u>**6**</u> | <u>**0**</u> |
| 925 | <u>**0**</u> | *17* | <u>**11**</u> | <u>**0**</u> | <u>**10**</u> | <u>**10**</u> | <u>**14**</u> | <u>**0**</u> | <u>**12**</u> | <u>**0**</u> | *24* | *25* | *24* | <u>**0**</u> | <u>**11**</u> | <u>**0**</u> | <u>**0**</u> |
| 1083 | *19* | *18* | *20* | *12* | *20* | *29* | *31* | *21* | *22* | *25* | *27* | *20* | *23* | *25* | *27* | *24* | *22* |

**Table 3:**
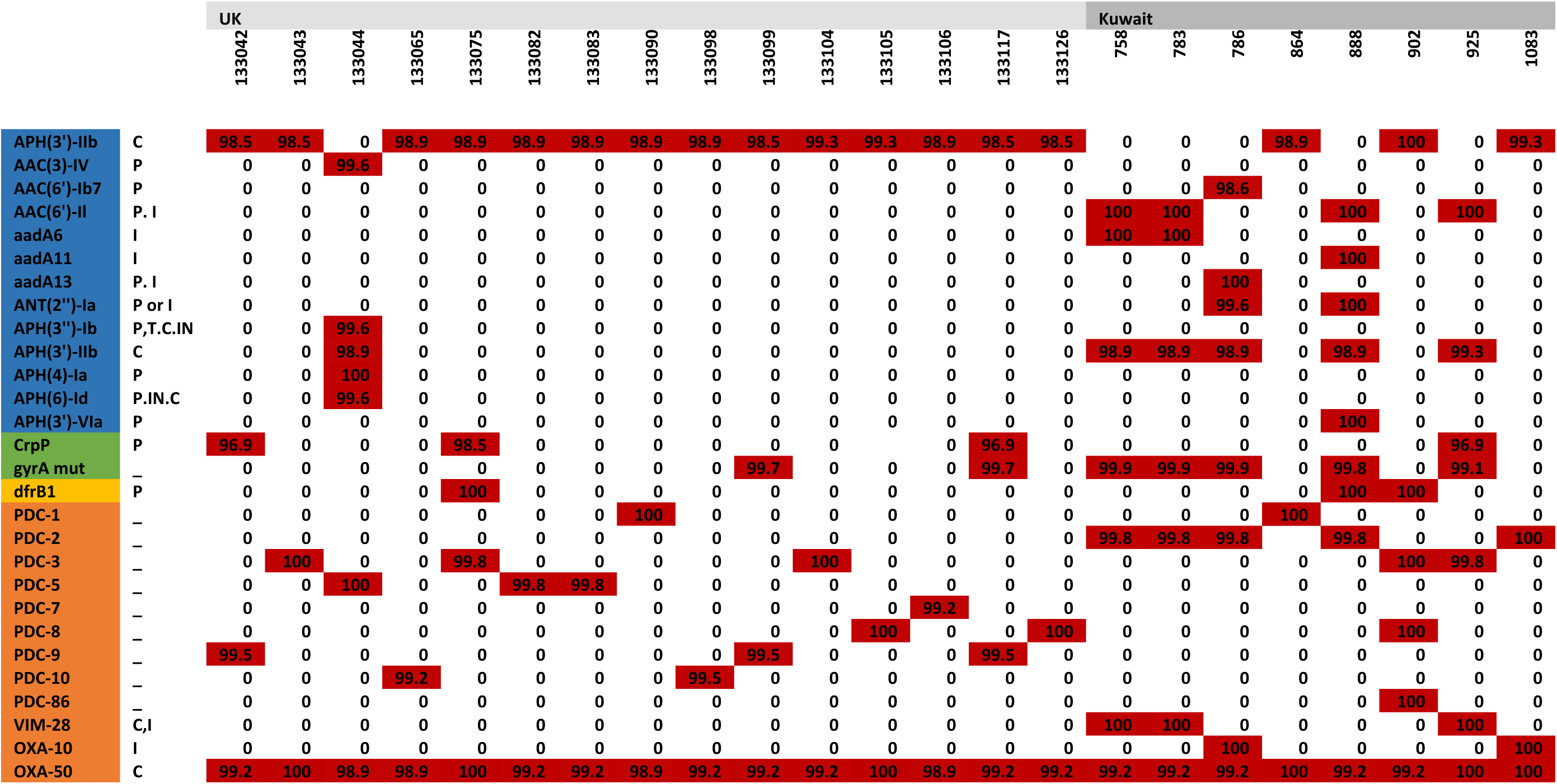
Antimicrobial resistance genes identified in *P. aeruginosa* UTI isolates through whole genome sequencing and analysis using the CARD database. Percentage sequence identity is reported. These genes are involved in promoting resistance to multiple of classes of antibiotics such as aminoglycosides (Blue), Fluoroquinolones (green), trimethoprim (yellow) and β-lactamases (orange) in UK & Kuwait isolates with resistance induced by C= Chromosome, I=Integrons=Transposons, IN=Integrative element.

The highest prevalence of resistance was to ciprofloxacin, a fluoroquinolone antibiotic utilised to treat *P. aeruginosa* infections. 10/14 isolates were resistance to this antibiotic with the remainder deemed susceptible with increased exposure. For four isolates, there was no zone of inhibition to ciprofloxacin and three of these also showed complete resistance to another fluoroquinolone, levofloxacin.

Resistance in the UK isolates was compared to a limited collection of resistant isolates from Kuwait. In contrast to the UK isolates, the isolates from Kuwait displayed very high levels of resistance (Table 2), albeit there appear to be bias in the selection of the panel. Seven out of eight isolates studied displayed multidrug resistance. Two isolates (758 and 783) were resistant to all antibiotics for which breakpoints were available. A broth microdilution MIC was performed to determine whether isolate 758 and 783 was resistant to the polymyxin colistin. Colistin was effective to both isolates at 2 mg/L, indicating that it might be utilised as a last-resort antibiotic to treat these strains. These isolates must be regarded as having a significant potential for pan resistance, since the obtained value of 2 mg/L indicates the threshold for classification as sensitive (sensitive ≥2, resistant <2).

Isolate 786 was only susceptible (increased exposure) to imipenem and meropenem. The other isolates displayed resistance to many classes on antibiotics. Isolate 1083 was the least resistant with resistance only to cefepime and ciprofloxacin.

### Genomic DNA analysis

To investigate the genomic context of UTI isolates in this investigation, whole genome sequencing was used to the core SNP phylogeny of isolates from the United Kingdom and Kuwait. Unusually, group 2 (PA14-like) isolates were more prevalent in the clinical UTI isolates. This contrasts with the prevalence of group 1 (PAO1-like) isolates within the Pseudomonas.com database (40). However, due to the small size of the panel, the results may not be representative of the wider *P. aeruginosa* clinical UTI population. To demonstrate the distribution of these UTI isolates amongst the wider population, a phylogenetic tree was created using sequenced isolates from KOS *et al*., (2015) (Figure 5) (41). Three Kuwaiti isolates clustered on the same branch of the phylogenetic tree, indicating that they are closely related.

**Figure 5:**
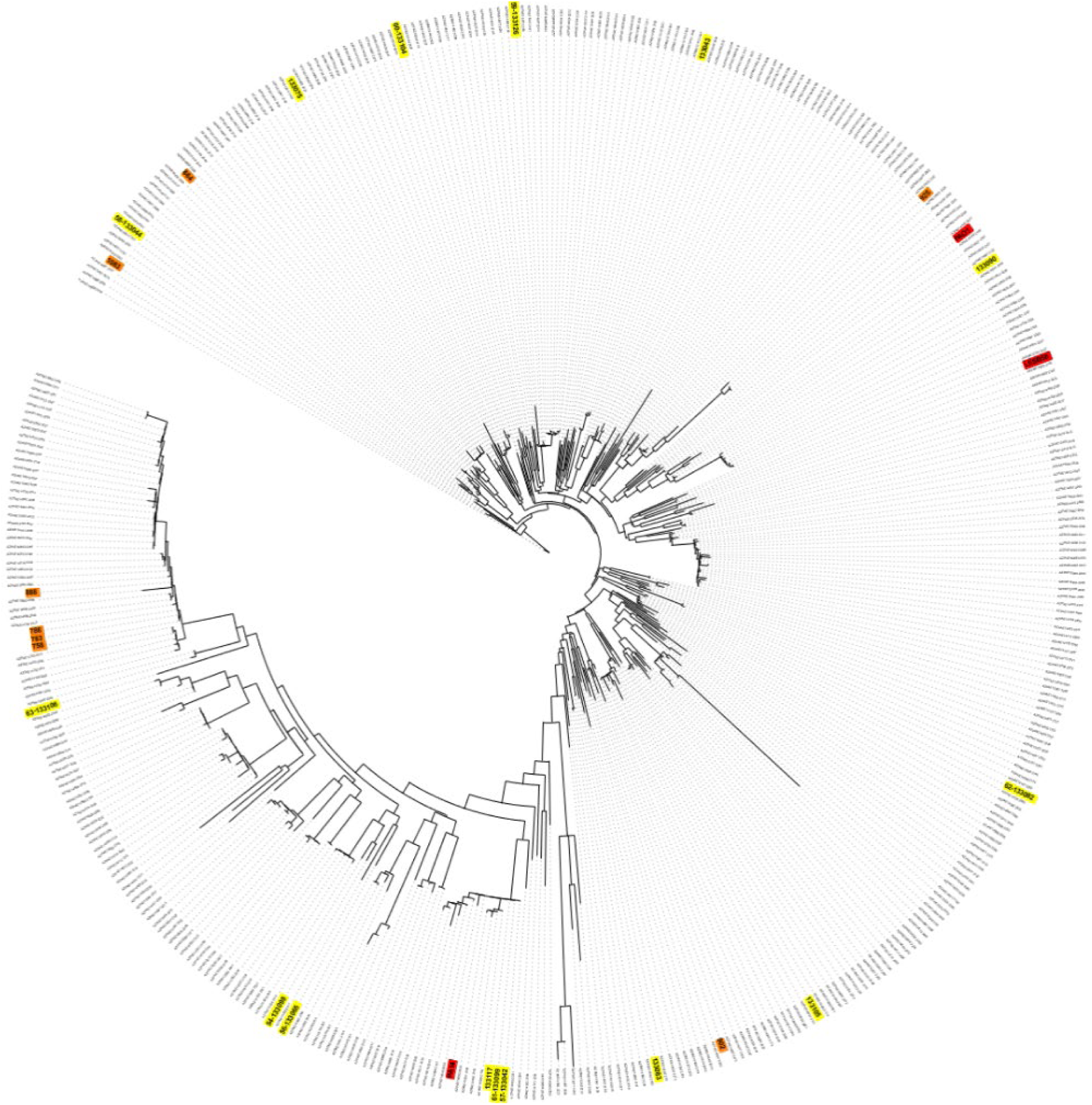
Phylogenetic tree of isolates from the United Kingdom and Kuwait based on their genomic composition, based on a core SNP phylogenetic tree using sequenced *P. aeruginosa* isolates from Principal strains (41) are color-coded in red; UK isolates in yellow, and Kuwaiti isolates in orange.

### Carriage of resistance genes in UTI isolates

CARD was used to identify resistance-associated genes in the UK UTI isolates. AAC(3)-IV was present only in isolate 133044. The gene encodes an aminoglycoside 3-N-acetyltransferase enzyme, which inactivates aminoglycosides including gentamicin and tobramycin, through enzymatic acetylation. Isolate 13044 displayed resistance to aminoglycosides. Further genes associated with aminoglycoside resistance were also identified including APH(3’)-Ib, APH(4)-Ia (both plasmid-encoded aminoglycoside phosphotransferase in *E. coli*), APH(6)-Id (a mobile genetic element-encoded aminoglycoside gene)(42) and APH(3’)-IIb, a chromosomal-encoded aminoglycoside(43).

Three isolates carried the gene crpP(44), which was linked to ciprofloxacin resistance however, this has been questioned and only 1/3 of these isolates were resistant to ciprofloxacin(45). In contrast, a gyrA mutation was identified in two isolates (133117 and 133099) and this corresponded to high-level resistance to both ciprofloxacin and levofloxacin. dfrB1 was identified in one isolate (133065) and encodes a plasmid-associated trimethoprim-resistant dihydrofolate reductase. A PDC variant of extended-spectrum beta-lactamase was identified in all isolates (46).

The isolates from Kuwait were highly resistant. AAC(6’) family genes of aminoglycoside 6’-N-acetyltransferase enzymes which inactivate gentamicin and tobramycin through enzymatic acetylation, were identified in 50% of the isolates with 2 isolates carrying AAC(6’)-Ib7 is a plasmid-encoded aminoglycoside acetyltransferase in *Enterobacter cloacae* and *Citrobacter freundii*.

Particularly resistant to aminoglycosides were isolates from Kuwait, with XDR isolates 758 and 783 being the most resistant in both panels (Table 7). These isolates contained the genes for the aminoglycoside transferase enzyme AAC (6’)-ii and the aminoglycoside adenylyltransferase aaA61, which both enzymatically inactivate aminoglycoside antibiotics. In addition, both isolates contained the plasmid-encoded *E. coli* phosphotransferase APH(3’)-Ib. This gene was detected in additional Kuwaiti isolates (786, 888, and 925) with similar aminoglycoside resistance. Isolate 888 contained five aminoglycoside resistance genes: AAC (6’)-ii, aadA11, ANT(2”)-Ia, APH(3’)-Ib, and APH(3’)-Via. Only isolate 133044 in the UK panel contained the aminoglycoside resistance genes APH(3”), APH(4), APH(3’)-Iib, and APH(6)-Id. This result demonstrates a correlation with the resistance patterns uncovered by the disc diffusion assay. Isolates 133042, 133075, 133117, and 925 all contained quinolone resistance genes such as *crpP*. Chávez-Jacobo *et al.* (2018) (44) described the enzyme product CrpP as a novel ciprofloxacin-modifying agent. Only one isolate, 133117 (crpP+), exhibited ciprofloxacin resistance in disc diffusion testing. Isolates from Kuwait had the highest frequency (62.5%) of mutations in gyrA that cause fluoroquinolone resistance.

Genes encoding β-lactamases are detected in all UK and Kuwaiti isolates. OXA-50 is encoded chromosomally and is present on all isolates. The other identified OXA type was OXA-10, a resistance gene detected in only one isolate (786). *P. aeruginosa* contains class C AMR genes encoding PDC - lactamases (46). All of the isolates contained at least one PDC enzyme, although the types varied. PDC-1, PDC-2, PDC-3, PDC-4, PDC-5, PDC-7, PDC-8, PDC-9, PDV-10, and PDC-86 were detected. PDC-2 is present in 26% of all tested UTI isolates, followed by PDC-3, which was only detected in isolates 758,783,778 and 888 from Kuwait. PDC-5 and PDC-9 were unique to UK isolates, with the former detected in 133044, 133082, and 130083 and the latter in 133042, 133099, and 133117. The remaining isolates contained PDC-1 (133090, 864) as well as PDC-10 (133065, 133098), PDC-7 (133106), and PDC-86. (902).

A single Verona integron-encoded metallo-β-lactamase (VIM) type gene was identified in three isolates. The VIM-28 gene was detected in these *P. aeruginosa* isolates. Other identified resistance genes include *sul1*, which is acquired through horizontal gene transfer and incorporated into the bacterial genome (47). The gene was first identified in *E. coli* and confers sulphonamide antibiotic resistance. The gene is detected in 62% of the Kuwaiti isolates and was notably absent in the UK panel of isolates.

## Discussion

Understanding UTIs is of paramount importance as UTIs are one of the most common bacterial infections worldwide. These infections can cause severe discomfort and pain to those afflicted, leading to a reduced quality of life and if left untreated, UTIs can progress to serious conditions, such as urosepsis (48). Additionally, the overuse and misuse of antibiotics in UTI treatment contribute to antibiotic resistance, a global health crisis. *P. aeruginosa* is largely understudied in the UTI context but can result in complex and resistant infections. By gaining an insight into phenotypic and genotypic variation, effective treatment strategies, preventive measures, and insights into the broader issue of antibiotic resistance may be gained.

*P. aeruginosa* can cause both acute and chronic infections. Chronicity is often associated with decreased growth under laboratory conditions. Here, the behaviour and fitness of UTI isolates was established by studying their growth under laboratory conditions. The majority of UTI isolates (71%) displayed a decreased AUC. Despite this, the generation time was relatively rapid with the majority of isolates showing faster generation than PAO1. Two isolates had a much slower generation time. This may reflect long term adaptation to the urinary environment. However, no information is available regarding the length of infection and therefore this cannot be confirmed. Overall, the capacity of UTI isolates to thrive in the laboratory environment is heterogeneous although many isolates display fast growth dynamics akin to environmental or acute clinical isolates.

*P. aeruginosa* displays an excellent capacity to form biofilms on both abiotic and biotic surfaces. All the UTI isolates showed some degree of biofilm ability however 79% displayed significantly decreased biofilm compared to PAO1. Multiple studies have documented the ability of *P. aeruginosa* isolated from multiple infection sites to form biofilms (49,50). *P. aeruginosa* can cause middle ear infections in cholesteatoma patients (51,52). 83% of otitis media *P. aeruginosa* isolates are efficient biofilm producers, with substantially higher levels of biomass produced than PAO1. The capacity to form biofilms may contribute to persistent infections in cholesteatoma patients (53). Similar to the UTI isolates in this study, PAHM4, an isolate from a person with non-CF bronchiectasis, produced substantially less biofilm than PAO1 (54). A study of 101 keratitis isolates revealed that isolates produced between 17% and 242% of the PAO1 control and that the ability to form biofilms was associated with poorer clinical outcomes (55). Schaber *et al.* (2007) compared 5 QS-deficient isolates from respiratory, cutaneous, and UTI infections to PAO1 to evaluate biofilm formation. The UTI isolate CI-5 was obtained from an 82-year-old patient who developed sepsis due to CAUTI. CV staining and comparison of all isolates revealed that, among all QS-deficient isolates, the UTI isolate produced the highest biomass and up to 82% capacity of PAO1 (56). A study comparing the capacity of uropathogenic *P. aeruginosa* serotypes to form biofilms *in vitro* using CV staining revealed variation between various serotypes. In particular, O11 serotypes produced the greatest biofilms and were more likely to be antibiotic-resistant, in contrast, O6 strains produced the weakest biofilms (57). Vipin *et al.* (2019) evaluated the ability of CAUTI *P. aeruginosa* to produce biofilms; all isolates formed biofilms between 0.20 and 1.11 (OD580), indicating the diversity of these clinical isolates (58). Additionally, Tielen *et al*. (2011) conducted CV staining on 12 mid-stream urine isolates and 18 CAUTI isolates and found the latter more proficient at producing biofilms. The majority of isolates were found to be intermediate biofilm producers (59). Our results suggest that mid-stream UTI are not as proficient in forming biofilms as keratitis, otitis media and CAUTI isolates. The ability to form biofilms is heterogenous in the tested panel.

*P. aeruginosa* employs motility as a prominent virulence factor to aid infection in multiple niches (60). Tielen *et al*. (2011) reported that 100% of 12 UTI isolates were capable of swimming, in comparison to 16 out 18 CAUTI (88.8%). Loss of swimming function might be related to evolutionary adaptation mechanisms in chronic infections similar to those observed in the lungs of CF patients.(59). More than 95% of CAUTI isolates have the functional ability to swim according to Ruzicka, *et al*.(2014) (61).In this study, 80% of the UTI isolates exhibited the ability to swim. Swimming motility can allow bacteria to colonise new niches and initiates attachment and adherence on surfaces, contributing to biofilm formation including on catheter surfaces (62). Type IV pili facilitate twitching motility, which is used to traverse solid and semisolid surfaces (63). 75% of the 139 *P. aeruginosa* UTI isolates examined by Olejnickova *et al.* (2014) for twitching motility were found to be capable of twitching (61). Winstanley *et al.* (2005) examined the twitching motility of 63 keratitis isolates; 90% of the isolates demonstrated the capacity to twitch *in vitro* (64). In addition, mutants lacking twitching motility were incapable of colonizing the cornea (65). In our study, 86.6% of the isolates were able to twitch, indicating that this is a common characteristic of UTIs. Furthermore, 86% of UTI isolates tested in this study, exhibited a high level of swarming motility. Hyper-motile strains that swarm tend to create flat biofilms (66). There have been reports of up to 95% of *P. aeruginosa* CAUTI isolates with swarming motility (61). Interestingly, swarming is a key characteristic of *Proteus mirabils*, another urinary pathogen (67). This may indicate that swarming is advantageous in the urinary environment.

The WHO has identified carbapenem-resistant *P. aeruginosa* as a critical issue requiring intervention [36]. Specific information on AMR of *P. aeruginosa* UTI isolates in both the UK and Kuwait are limited (26,68,69). Phenotypic resistance screening revealed a relatively low level of resistance in isolates from the UK. However, pan resistance was identified in some isolates from Kuwait. This could suggest that Kuwait may be a hotspot for antibiotic resistance however, this small number of isolates were stored based on higher resistance and therefore represents a biased subset. A recent study in a major Kuwaiti hospital revealed that 32.1% of *P. aeruginosa* isolates were MDR, the majority of these were derived from urine culture (70). Extensive prospective studies should be conducted since most of the obtained AMR data in Kuwait are based on retrospective studies. Lack of surveillance programs, inadequate use of antibiotics, travel, and climate change all contribute to the spread of antimicrobial resistance in this region (19). Whole genome sequencing revealed resistance determinants in isolates from both geographical regions however, uncommon resistance genes were identified in the genomes of isolates from Kuwait. The Verone integron-encoded metallo-beta-lactamase (VIM) family are able to inactivate bet-lactam antibiotics and the unusual VIM-28 was identified in 4 isolates. VIM-28 is a metallo-β-lactamase initially reported in Egypt, the structure of the VIM-28 gene contains an uncommon integron configuration, with the gene cassette located downstream of the “*intI1*” gene. Egyptian nationals constitute one of Kuwait’s main ethnic minorities (71,72). Travelling between the two nations may help explain the presence of ^bla-^VIM-28 in both regions. In addition to the intrinsic OXA-50 Class D β-lactamase, which is present in all isolates, the OXA-10 Class D β-lactamase was also detected is isolate 786 and 1083. To the best of our knowledge, this is the first report of OXA-10 in Kuwaiti bacterial isolates.

Multiple isolates exhibited resistance to aminglycosides. Uncommon genes in *P. aeruginosa* including aadA6, aadA11 and aadA13 were observed. AadA-type genes encode aminoglycoside nucleotidyltransferases and these are often encoded on mobile genetic elements such as plasmids and integrons. Genes for further aminoglycoside-modifying enzymes were also observed including AAC (6’)-ii, ANT(2’’)-Ia and APH(3’)-Ib. Fluoroquinolone resistance was also observed. Mutations in *gyrA* and plasmid-associated CrpP enzyme would contribute to ciprofloxacin and levofloxacin resistance in these isolates (44). dfrB1 is a plasmid-associated, trimethoprim-resistant dihydrofolate reductase that was initially identified in *Bordetella bronchispetica* bacteria (73), suggesting that it may have been horizontally transmitted to *P. aeruginosa*. Due to the extended resistance and identified genes, it is possible that these isolates contain MDR plasmids. Recent research on Thai isolates revealed that *P. aeruginosa* carries MDR megaplasmids (74). Additional analysis, particularly using long read sequencing data, would reveal further information on plasmid carriage. The implications of these findings are alarming and suggest extensive AMR surveillance and management programmes would be beneficial.

In this study, only two MDR-isolates were detected in isolates originating from the United Kingdom, indicating that they were largely susceptible to antibiotics. In general, the prevalence of AMR *P. aeruginosa* uropathogens is regarded as low (18). This could be attributed to stricter antibiotic stewardship policies and increased public awareness of antibiotic misuse (75). According to Ironmonger *et al.*, a 4-year surveillance investigation detected only 45 non-susceptible *P. aeruginosa* uropathogens to carbapenems among 6985 isolates from the Midlands region of England (18).

The findings in this study reveal the genomic and phenotypic diversity of *P. aeruginosa* isolates from UTIs. The data indicate that UTIs can be caused by a wide variety of phenotypic and genotypic combinations. The success of a given infection is therefore likely a complex interplay between bacterial characteristics and host factors, such as hospitalisation and long-term catheter use. Sequencing conducted as part of this study detected the presence of antimicrobial resistance genes that could promote resistance to every class of antibiotics. Uncommon resistance genes were highly prevalent in the isolates from Kuwait, and this correlated with extremely high phenotypic resistance. However, the basis on which these isolates were selected is unclear and likely biased. Thus, additional study would need to determine the prevalence of such strains. In addition, a deeper understanding of how *P. aeruginosa* responds to the urinary environment would allow for a more in-depth examination of the bacterial pathogenesis and aid in the identification of novel therapeutic interventions.

## Author Notes

Accession numbers TBC.

## Abbreviations

AMR: Antimicrobial resistance
CARD: Comprehensive Antibiotic Resistance Database
CV: Crystal violet
CAUTI: Catheter associated infections
CF: Cystic fibrosis
CGR: Centre for Genomic Research
*E. coli*: *Escherichia coli*
EUCAST: European Committee on Antimicrobial Susceptibility Testing
GCC: Gulf Corporation Council
MIC: Minimum inhibitory concentrations
*P. aeruginosa*: *Pseudomonas aeruginosa*
UTI: Urinary tract infection
VIM: Verona integron-encoded metallo-β-lactamases
WGS: Whole Genome Sequencing
WHO: World Health organization
XDR: extensively drug resistant.

## Author statements

### Author contributions

HE., funding, investigation, formal analysis, writing – original draft preparation; R.F., conceptualisation, writing – review and editing, supervision, project administration, analysis; SH., Bioinformatic analysis; MM., Bioinformatic analysis; AD., Consultation and providing I Kuwait isolates; J.F., conceptualisation, writing – review and editing, supervision, project administration, analysis.

### Author Statement

We would like to thank CGR at the University of Liverpool for providing sequencing services.

